# Selection maintains protein interactome resilience in the long-term evolution experiment with *Escherichia coli*

**DOI:** 10.1101/2021.01.20.427477

**Authors:** Rohan Maddamsetti

**Affiliations:** Department of Biomedical Engineering, Duke University, Durham NC

## Abstract

Most cellular functions are carried out by a dynamic network of interacting proteins. An open question is whether the network properties of protein interactomes represent phenotypes under natural selection. One proposal is that protein interactomes have evolved to be resilient, such that they tend to maintain connectivity when proteins are removed from the network. This hypothesis predicts that interactome resilience should be maintained by natural selection during long-term experimental evolution. I tested this prediction by modeling the evolution of protein-protein interaction (PPI) networks in Lenski’s long-term evolution experiment with *Escherichia coli* (LTEE). In this test, I removed proteins affected by nonsense, insertion, deletion, and transposon mutations in evolved LTEE strains, and measured the resilience of the resulting networks. I compared the rate of change of network resilience in each LTEE population to the rate of change of network resilience for corresponding randomized networks. The evolved PPI networks are significantly more resilient than networks in which random proteins have been deleted.

Moreover, the evolved networks are generally more resilient than networks in which the random deletion of proteins was restricted to those disrupted in LTEE. These results suggest that evolution in the LTEE has favored PPI networks that are, on average, more resilient than expected from the genetic variation across the evolved strains. My findings therefore support the hypothesis that selection maintains protein interactome resilience over evolutionary time.

**Significance Statement:** Understanding how protein-protein interaction (PPI) networks evolve is a central goal of evolutionary systems biology. One property that has been hypothesized to be important for PPI network evolution is resilience, which means that networks tend to maintain connectivity even after many nodes (proteins in this case) have been removed. This hypothesis predicts that PPI network resilience should be maintained during long-term experimental evolution. Consistent with this prediction, I found that the PPI networks that evolved over 50,000 generations of Lenski’s long-term evolution experiment with *E. coli* are more resilient than expected by chance.

## Introduction

When redundant nodes and connections in a network carry a cost, removing those redundancies may increase efficiency at the expense of reducing resilience to unexpected disruptions. After a critical point, called the percolation threshold, further pruning can cause a catastrophic breakdown of connectivity and function (Callaway, et al. 2000). In the context of protein-protein interaction (PPI) networks, efficiency may refer to the cost of protein production (Kafri, et al. 2016).

Zitnik and colleagues (2019) formally defined network resilience to measure how quickly a network breaks down as nodes are removed. They then studied the evolution of PPI networks (also called protein interactomes) across the tree of life, concluding with the hypothesis that interactome resilience is favored during evolution. While interesting, this hypothesis is rather vague. To better understand the relevance of network resilience to evolutionary biology, we need to ask whether network resilience has any relevance or predictive power in additional contexts. If network resilience is a *necessary* property of evolved protein-protein interaction networks, then it should be maintained by selection during long-term evolution experiments. Here, I use the methods developed by Zitnik et al. (2019) to test this prediction, by examining how protein interactome resilience has evolved in Lenski’s long-term evolution experiment with *Escherichia coli* (LTEE).

In the LTEE, 12 initially identical populations of *E. coli* have evolved for more than 50,000 generations (Lenski, et al. 1991; Lenski 2017). The LTEE populations are named by the presence of a neutral phenotypic marker: populations Ara+1 to Ara+6 grow on arabinose, while populations Ara−1 to Ara−6 cannot (Lenski, et al. 1991). Many populations have lost unnecessary metabolic traits (Leiby and Marx 2014; Grant, et al. 2020), and many genes have been disrupted by loss-of-function mutations, in part caused by the evolution of elevated mutation rates in several populations (Tenaillon, et al. 2016; Couce, et al. 2017; Good, et al. 2017; Maddamsetti and Grant 2020a).

Despite evidence for genomic and phenotypic streamlining, it is unknown how PPI network resilience has evolved during the LTEE. To examine this question, I compared the rate of change of protein interactome resilience in each LTEE population to the expected rate of change in corresponding sets of randomized networks (Material and Methods). I compare rates of change of interactome resilience between real and simulated networks, because this approach is simple and accounts for phylogenetic correlations among genomes from the same population; it treats the independent LTEE populations as the appropriate unit of statistical replication. Importantly, the randomized networks within each population have no such phylogenetic correlations, in order to reduce the computational cost of sampling large numbers of statistically independent replicates. For robustness, I analyzed two curated datasets of protein-protein interactions in *E. coli*, which I refer to as Zitnik interactome (Zitnik, et al. 2019) and the Cong interactome (Cong, et al. 2019). Overall, protein interactome resilience is higher in the LTEE than in the randomized networks, indicating that this system-level property is being maintained over long-term experimental evolution.

## Material and Methods

### Datasets

I downloaded a table of nonsense SNPs, small indels, mobile element insertions, and large deletions affecting protein-coding regions of 264 genomes of LTEE clones isolated at 11 timepoints through 50,000 generations (Tenaillon, et al. 2016) using the web application at: https://barricklab.org/shiny/LTEE-Ecoli. The underlying data are also available at: https://github.com/barricklab/LTEE-Ecoli. Because these mutations disrupt protein reading frames, I use them as a proxy for loss-of-function mutations in the LTEE. For this reason, I call these types of mutations “gene disruptions”, and call genes that are affected by these types of mutations “disrupted genes” for short. In the analyses restricted to 50,000 generation LTEE clones, I used single clones from each of the 12 populations (the LTEE 50,000 generation ‘A’ clones) to maximize statistical independence.

I used two different PPI datasets for *E. coli*. First, I used the curated PPI set published by Zitnik et al. (2019). This set of interactions corresponds to the interactome for species 511145 in their cross-species interactome dataset. For robustness, I also used a second *E. coli* protein interactome that has been published by Cong and colleagues (2019). This set of interactions includes those from protein structures in the Protein Data Bank (their Table S4), the Ecocyc database (their Table S5), yeast two-hybrid experiments (their Table S6), affinity purification and mass spectrometry (their Table S7), and those supported by known and high-confidence novel prediction coevolutionary information (their Tables S8 and S10, respectively). The Zitnik and Cong interactomes were filtered based on the genes they share with the *E. coli* B str. REL606 genome (the ancestral LTEE clone).

A list of essential and nearly essential genes in the LTEE ancestral clone REL606 was taken from Supplementary Table 1 of Couce et al. (2017), who identified these genes through transposon mutagenesis and sequencing. A comparison to the 57 genes with clear evidence of parallel evolution (i.e. two or more nonsynonymous mutations) in nonmutator lineages of the LTEE (Tenaillon, et al. 2016; Maddamsetti, et al. 2017) uses the data reported in supplementary table 2, Supplementary Material online of Tenaillon et al. (2016).

### Network resilience analysis

The snap.py python module interface to the Stanford Network Analysis Platform (Leskovec and Sosič 2016) was used to generate a graph representing the ancestral REL606 interactome. Then, a protein interactome network was generated for each LTEE genome, by pruning the REL606 interactome of nodes (proteins) and edges (interactions) affected by gene disruptions in the given genome. Network resilience was calculated using the method described in Zitnik et al. (2019). In brief, resilience measures how well a network resists fragmentation into many small, isolated components as increasing fractions of nodes are randomly removed. Specifically, the entropy of the distribution of component sizes in the network is calculated, normalized by the logarithm of the total number of nodes in the original network. A fraction *f* of the nodes of the graph are randomly removed, ranging from 0.01 to 1.0, and the entropy of the network component distribution is calculated for each value of *f*. The entropy of the network component distribution increases monotonically as *f* increases. The area under the curve (AUC) of the mapping from *f* to the entropy of the network component distribution is calculated numerically, and network resilience is defined as 1 – AUC. This resilience calculation was conducted 100 times per genome. The average network resilience out of these 100 samples was used as the estimate for a genome’s network resilience for a given simulation run. 100 simulations of this entire algorithm were conducted, such that network resilience was calculated 10,000 times for each genome. The trend over time in each population was then plotted, and a linear regression was fit to estimate the change in network resilience over time in each LTEE population. The *y*-intercept was fixed as the mean estimate for the resilience of the ancestral LTEE clone REL606 over all 100 simulation runs, so that the linear regression only fits one parameter (a slope representing network resilience over time). I then constructed two null distributions for comparison to the actual data as follows:

#### Randomization over all genes in REL606

For each of the 264 LTEE genomes (Tenaillon, et al. 2016), I constructed a corresponding randomized interactome network (Figure 1). The number of disrupted genes (i.e. genes removed from the ancestral interactome network) was fixed to the number of nonsense SNPs, small indels, mobile element insertions, and large deletions affecting protein-coding regions in the given LTEE genome. Then, a selection of genes in the ancestral REL606 genome was drawn at random. These genes were removed from the ancestral interactome network, leaving a randomized network with the same number of disrupted genes as in the given evolved genome. The resilience of these randomized networks was calculated as described above.

**Figure 1.**
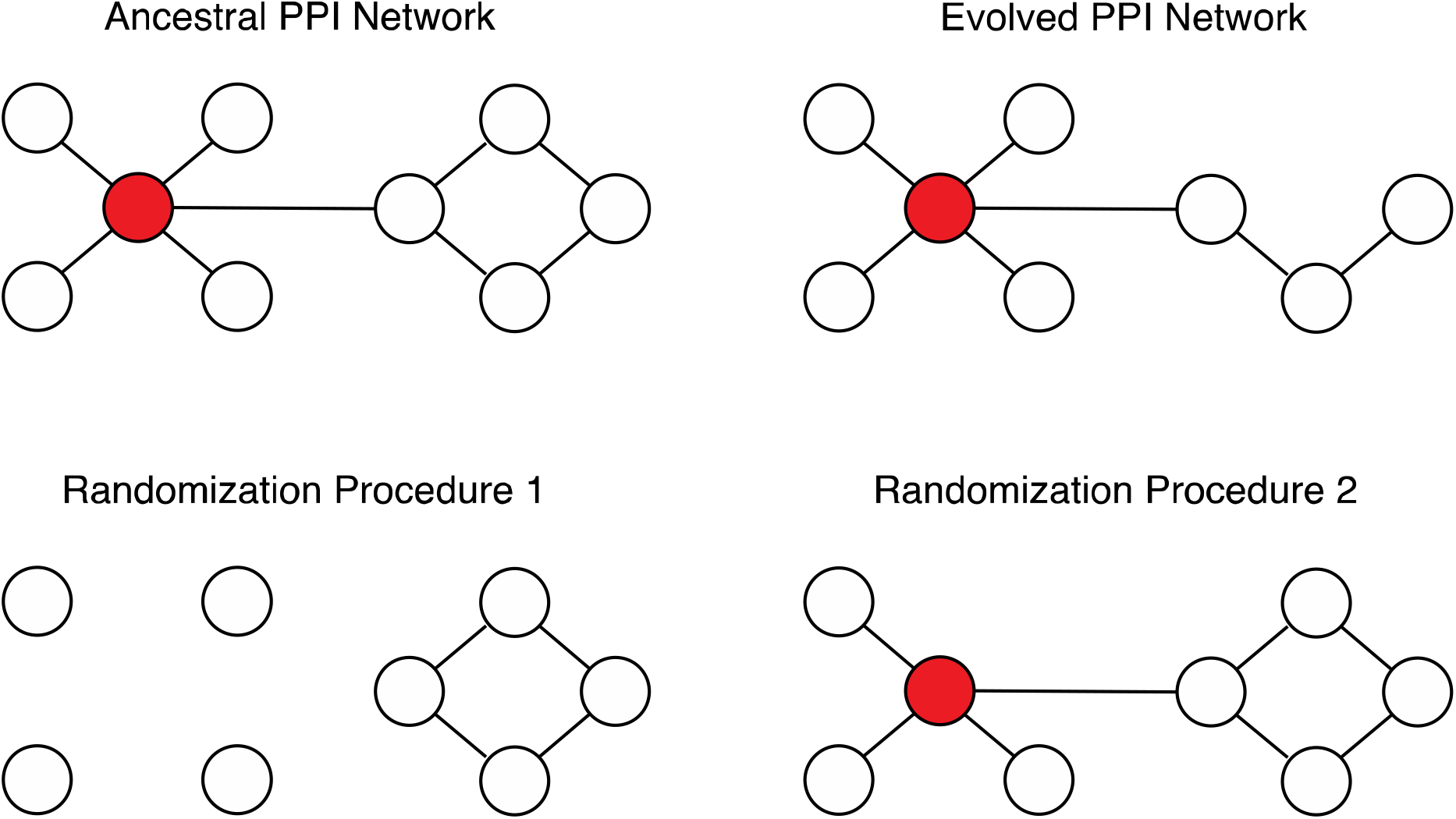
Illustration of ancestral, evolved, and randomized networks. Nodes represent proteins, and edges represent protein-protein interactions. Red nodes indicate essential proteins which cause lethal phenotypes if disrupted by nonsense SNPs, small indels, mobile element insertions, or large deletions. The PPI network of the ancestral bacterial clone is shown at top left. The PPI network of an evolved bacterial clone, in which one protein has been disrupted, is shown at top right. The randomization procedures sample subnetworks of the ancestral PPI network. Each randomized network corresponds to an evolved PPI network: the number of proteins disrupted in the randomized network is fixed to the number disrupted in the corresponding evolved PPI network. The first randomization procedure used in this paper (bottom left) samples all protein-coding genes in the ancestral clone for disruption, including those encoding essential proteins. The second randomization procedure used in this paper (bottom right) samples protein-coding genes for disruption based on the number of evolved populations that contain disruptions of that gene. See Material and Methods for further details.

#### Randomization over genes disrupted across LTEE populations

For each of the 264 LTEE genomes (Tenaillon, et al. 2016), I again constructed a corresponding randomized interactome network (Figure 1). As before, the number of disrupted genes was fixed to the number of nonsense SNPs, small indels, mobile element insertions, and large deletions affecting protein-coding regions in the given LTEE genome. In this case, however, a selection was drawn from the genes that were disrupted across all LTEE populations. I weighted the probability of selecting each gene based on the multiplicity of observed disruptions of that gene across LTEE populations. Specifically, the set of disrupted genes for each LTEE population was calculated, then the multiplicity of those genes across populations was calculated, such that a gene that was disrupted in 5 populations would be represented 5 times for sampling, while a gene that was never disrupted in any LTEE genomes would be omitted. The resilience of the resulting randomized network was calculated as described above.

The differences between the slopes for linear regressions for the actual data and the randomized data for each population were tabulated, and a Wilcoxon signed-rank test was used to test whether the paired difference between slopes, over all LTEE populations, was significantly greater than zero.

### Single gene disruption analysis

Interactome resilience was calculated for the ancestral LTEE clone and the 12 50,000 generation LTEE ‘A’ clones, representing all 12 LTEE populations. For each clone, and for every protein-coding gene, the given protein was removed from the interactome network of the given clone, and interactome resilience was re-calculated. If the given protein did not have any interactions in the network, then the resilience of the resulting network was set to the resilience of the original network for the clone. As before, each resilience calculation was conducted 100 times per genome, and the average network resilience out of these 100 samples was used to estimate the clone’s network resilience. However, only one simulation of this entire algorithm was conducted, such that network resilience was calculated 100 times for each of the ~4,000 networks corresponding to single-gene disruptions covering all protein-coding genes for each of the 13 genomes (ancestor and 12 clones) analyzed.

## Results

### LTEE PPI networks are more resilient than PPI networks with random proteins deleted

I calculated the resilience of randomized counterparts of the evolved LTEE PPI networks, in which proteins to delete from the network were sampled at random. In this randomization scheme, a protein may be removed from the PPI network, regardless of its essentiality in *E. coli* (Figure 1). For this reason, I expected that the evolved LTEE PPI networks would be more resilient than the randomized PPI networks, and this was indeed the case (Zitnik interactome: Wilcoxon signed-rank exact test *p* = 0.00024; Cong interactome: Wilcoxon signed-rank exact test *p* = 0.00024). This result can be seen by comparing the red and yellow slopes in Figure 2.

**Figure 2.**
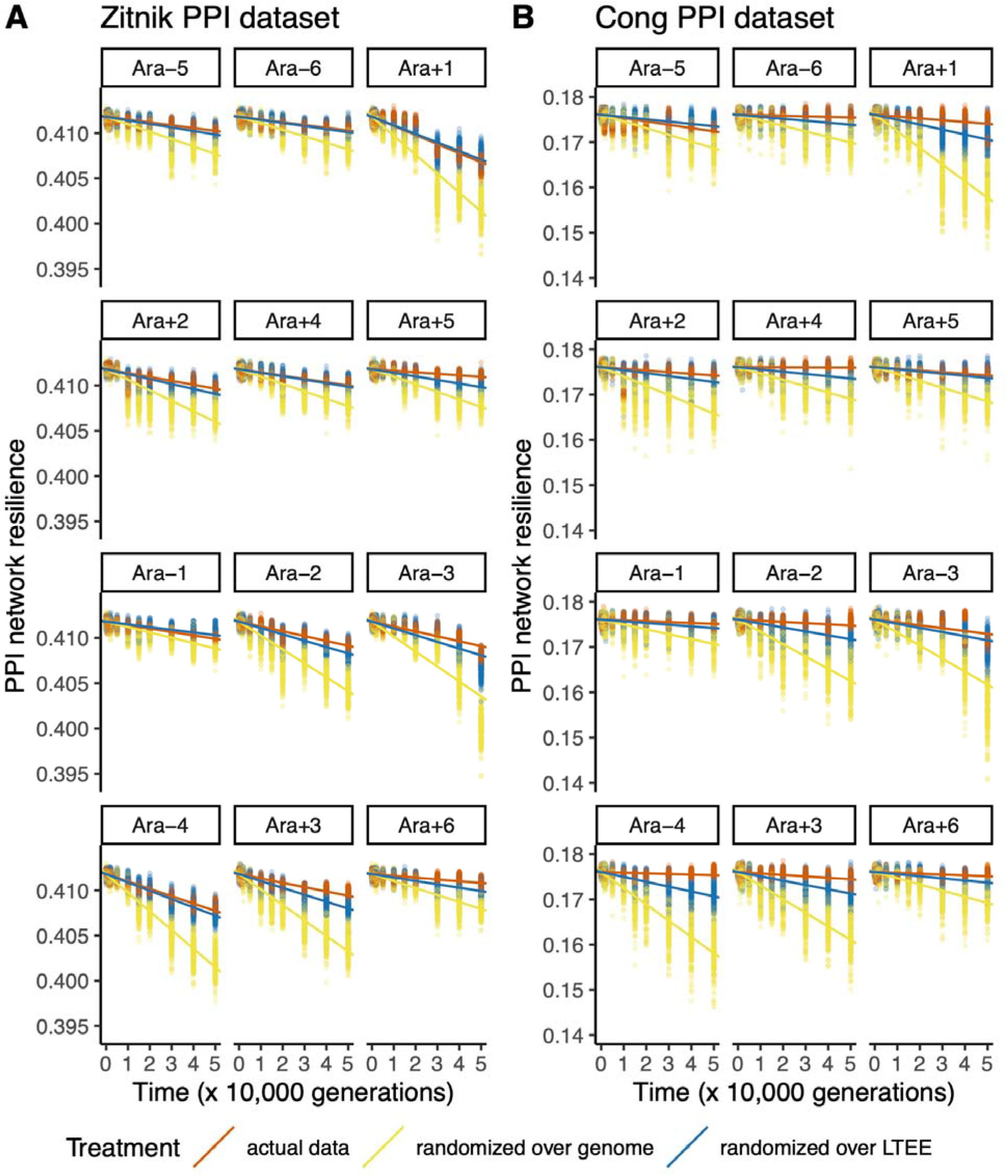
Protein-protein interaction (PPI) networks in the LTEE (red) lose network resilience more slowly than randomized networks generated using either all genes in the REL606 genome (yellow) or only those disrupted in the LTEE (blue). The top six populations have the ancestral point-mutation rate, while the bottom six populations evolved elevated point-mutation rates. A) Analysis based on the *E. coli* PPI network published in Zitnik et al. (2019). B) Analysis based on the *E. coli* PPI network published in Cong et al. (2019).

The analysis in Figure 2 also shows that the Ara+1 population is an outlier: randomized Ara+1 networks are expected to have very low resilience, even though the Ara+1 population has maintained the wild-type point mutation rate (see Figure 2 legend). This finding can be explained by the high transposon insertion mutation rate that evolved in this population (Papadopoulos, et al. 1999; Maddamsetti and Grant 2020a; Consuegra, et al. 2021). The Ara+1 population has also lagged behind the others in mean fitness (Wiser, et al. 2013; Grant, et al. 2020; Consuegra, et al. 2021)

### LTEE PPI networks are more resilient than PPI networks with gene disruptions sampled across LTEE populations

In part, the previous finding could be caused by sampling unrealistic randomized networks. For instance, the previous method permits the sampling of randomized networks that lack essential ribosomal proteins, which is biologically implausible (Figure 1). I therefore conducted a second test, in which I restricted the proteins sampled for deletion to those that were disrupted in at least one LTEE population. Here, the probability of sampling proteins for deletion was weighted by the frequency of observed disruptions across LTEE populations (Figure 2). This resampling procedure takes parallel genetic evolution into account (Woods, et al. 2006; Ostrowski, et al. 2008; Tenaillon, et al. 2016), because proteins that are disrupted multiple times across populations are more likely to be sampled. I found that that evolved LTEE PPI networks, on the whole, are more resilient than randomized PPI networks generated from gene disruptions tabulated across all 12 LTEE populations (Zitnik interactome: Wilcoxon signed-rank exact test *p* = 0.0061; Cong interactome: Wilcoxon signed-rank exact test *p* = 0.0012). This result can be seen by comparing the red and blue slopes in Figure 2.

### Genes in large deletions in the LTEE neither show physical modularity nor fewer interactions

The resampling procedures operate on the level of individual gene disruptions and losses, such that large deletions are replaced by a sample of individual gene disruptions. Therefore, the resampling procedures could bias the results by breaking up the block structure of multi-gene deletions. This would matter if disrupting a block of *x* genes has less of an effect on PPI network resilience than disrupting *x* genes across the genome.

I examined two PPI properties through which the absence of large deletions in the randomized networks could have an effect. First, systematic bias could be introduced if genes within large deletions tend to have fewer interactions than genes affected by small indels or nonsense mutations. Second, systematic bias could be introduced if interactions within large deletions show physical modularity, such that genes within large deletions preferentially interact with each other, but not with genes elsewhere on the chromosome.

I find no difference in PPI degree between genes knocked out by multi-gene deletions, and those disrupted by single-gene mutations in the 50,000 generation LTEE clones (Zitnik interactome: Wilcoxon rank-sum test *p* = 0.46; Cong interactome: Wilcoxon rank-sum test *p* = 0.26). Furthermore, I find that interactions removed by multi-gene deletions in 50,000 generation LTEE clones are *further apart* in the genome, on average, than interactions removed by single-gene disruption mutations. For the case of the Zitnik interactome, interactions removed by multi-gene deletions have a mean genomic distance of 1,071,063 base-pairs, compared to a mean genomic distance of 1,054,403 base-pairs for interactions removed by single-gene disruptions (Wilcoxon rank-sum test: *p* = 0.0364). For the case of the Cong interactome, interactions removed by multi-gene deletions have a mean genomic distance of 714,316 base-pairs, compared to a mean genomic distance of 583,624 base-pairs for interactions removed by single-gene disruptions (Wilcoxon rank-sum test: *p* < 10^−4^). Therefore, the differences between realized and randomized PPI network resilience in the LTEE do not seem to be an artifact of the resampling procedure, at least with regard to PPI degree and the aspects of physical modularity that I examined.

### Purifying selection on essential genes in the LTEE ancestral clone is insufficient to explain the maintenance of network resilience in the LTEE

What evolutionary forces are responsible for maintaining protein interactome resilience in the LTEE? Interactome resilience is not a target of positive selection, because mean population fitness increases in each LTEE population (Wiser, et al. 2013) while interactome resilience decreases (Figure 2). Therefore, interactome resilience negatively correlates with fitness gains in the LTEE (Figure 3). The only remaining explanation is that network resilience is being maintained by purifying selection. Still, it is unclear whether protein interactome resilience is under direct selection, or whether its maintenance is a byproduct of purifying selection on correlated phenotypes.

**Figure 3.**
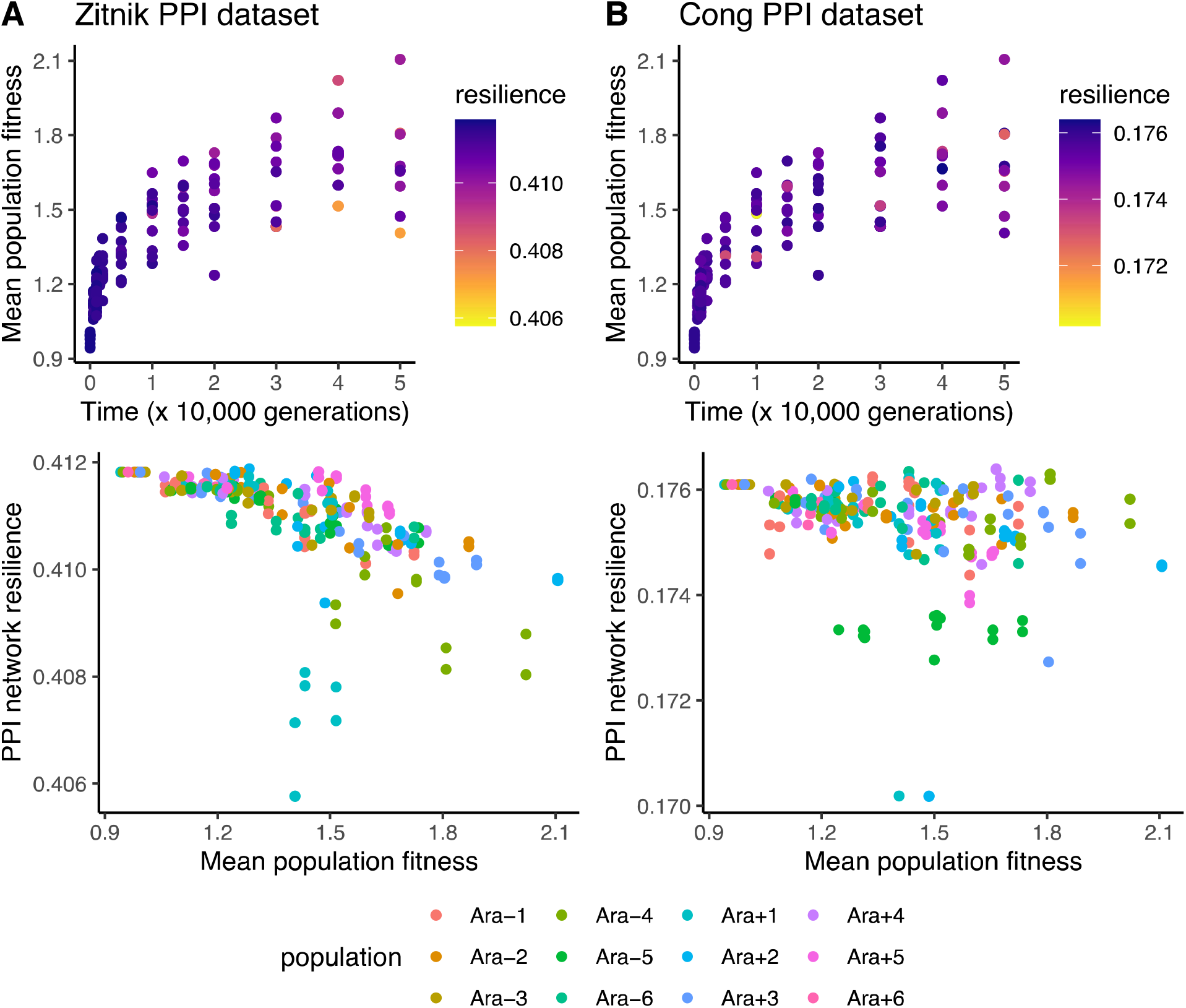
PPI network resilience negatively correlates with mean population fitness, measured by direct competition assays against reference LTEE clones (Wiser et al. 2013). The top panels show the fitness measurements from Wiser et al. (2013); the colors denote the PPI network resilience of genomes sampled from the corresponding populations and time points. The bottom panels show the negative correlations between mean population fitness and PPI network resilience. A) Analysis based on the *E. coli* PPI network published in Zitnik et al. (2019). The negative correlation between fitness and resilience is significant (Pearson’s product-moment correlation: *r* = −0.59, *p <* 10^−16^). B) Analysis based on the *E. coli* PPI network published in Cong et al. (2019). The negative correlation between fitness and resilience is significant (Pearson’s product-moment correlation: *r* = −0.27, *p <* 10^−4^).

As essential genes cause lethal phenotypes when disrupted, direct purifying selection against the disruption of essential genes could cause indirect purifying selection on interactome resilience. I hypothesized that disruptions of essential genes would have disproportionately negative effects on network resilience. While I found evidence of direct purifying selection on essential genes, I found limited evidence for the hypothesis that direct purifying selection on essential genes indirectly maintains interactome resilience.

I examined a set of 541 essential and nearly essential genes that were identified in REL606 (Couce et al. 2017). These genes are highly enriched for protein-protein interactions compared to the remaining 3,571 non-essential genes (one-sided Wilcoxon rank-sum test: *p <* 10^−58^ for Zitnik PPI dataset, *p <* 10^−10^ for Cong PPI dataset). Essential and nearly-essential genes also show a significant signal of purifying selection in the 50,000 generation LTEE clones: they contain 44 out of 941 gene disruptions in the 50,000-generation LTEE genomes, while the total length of essential and nearly-essential genes is 499,180 bp, out of 3,962,143 bp representing the total length of all protein-coding genes (one-sided binomial test: *p <* 10^−15^). It should be noted that the 23 of the 57 genes with clear evidence of parallel evolution (i.e. two or more nonsynonymous mutations) in nonmutator lineages of the LTEE (Tenaillon, et al. 2016; Maddamsetti, et al. 2017) are essential or nearly essential. This association between essentiality and positive selection in the LTEE is highly significant (Fisher’s exact test: *p <* 10^−6^).

I then simulated the disruption of every single gene in REL606 to measure their effects on network resilience. In contrast with my initial expectation, I find that the majority of single gene disruptions in REL606 *increase* network resilience (Figure 4). Furthermore, disruptions of essential and non-essential genes have qualitatively similar effects on network resilience (Figure 4). When I examine the effect of disrupting every single gene in the 50,000 generation LTEE clones on interactome resilience, the results are largely strain-specific. Again, the trend for essential genes is similar to the trend for non-essential genes (Supplementary Figure S1). Together, these findings suggest that purifying selection on essential genes is not sufficient to explain the maintenance of network resilience in the LTEE.

**Figure 4.**
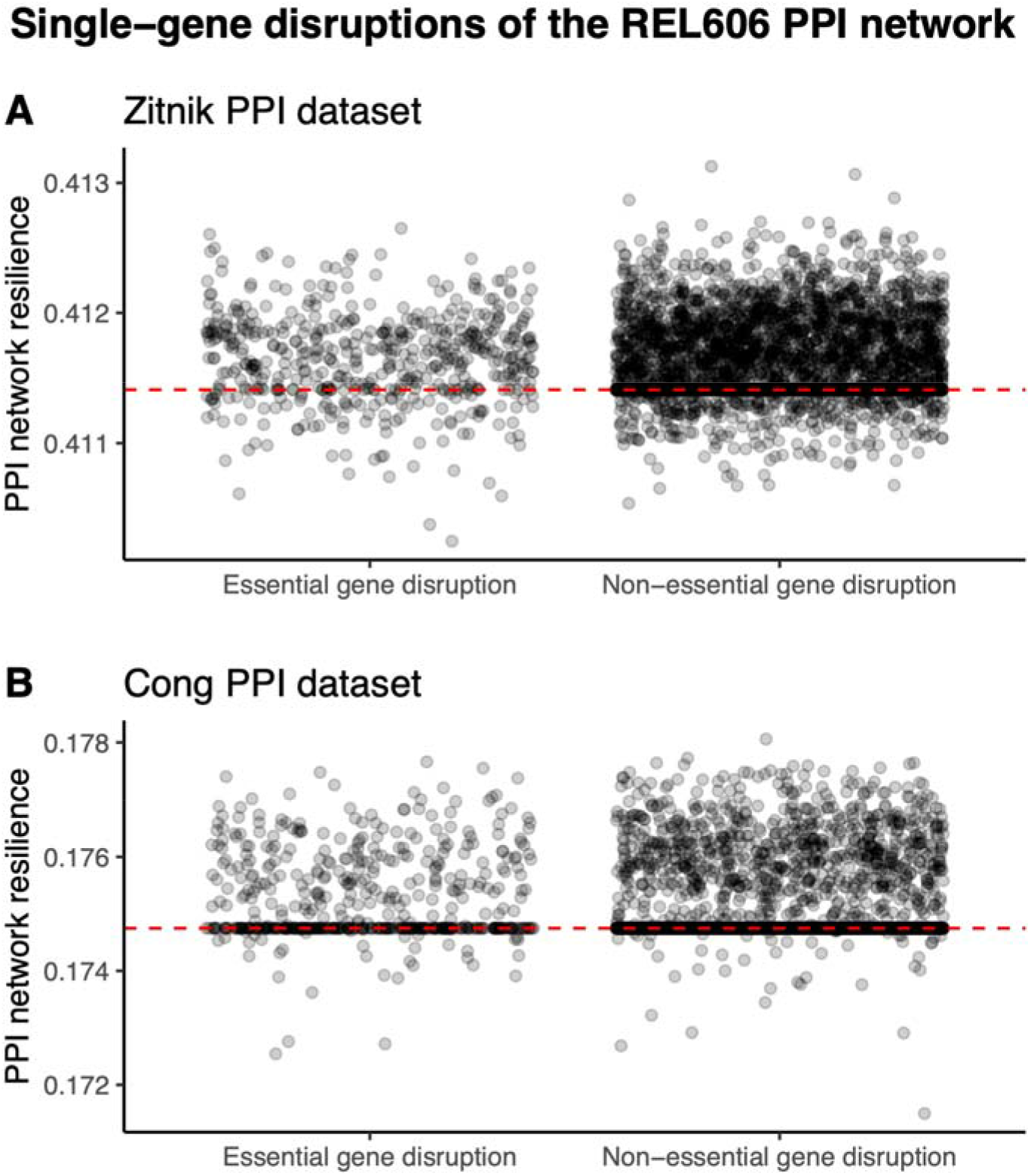
Single-gene disruptions tend to increase the resilience of the REL606 PPI network. Each point represents the resilience of the REL606 PPI network after a single gene has been removed. The distribution of resilience effects after removing essential and nearly essential genes is shown on the left, and the distribution of resilience effects after removing non-essential genes is shown on the right. The dashed red line indicates the resilience of the original REL606 network. A) Analysis based on the *E. coli* PPI network published in Zitnik et al. (2019). B) Analysis based on the *E. coli* PPI network published in Cong et al. (2019).

## Discussion

I find that evolved networks in the LTEE lose protein interactome resilience more slowly than expected, based on comparisons with networks with random gene disruptions and networks with gene disruptions weighted by their occurrence across LTEE populations (Figure 1). The second analysis controls for the biologically implausible PPI networks that would be created by sampling essential genes for disruption. Together, these results are consistent with Zitnik and colleagues’ general hypotheses that network resilience 1) is a genuine property of evolved protein-protein interaction networks and 2) is relevant for understanding how protein-protein interaction networks evolve.

Selection must be driving the maintenance of protein interactome resilience in the LTEE. Since protein interactome resilience negatively correlates with fitness gains in the LTEE (Figure 3), positive selection can be ruled out as a cause. Therefore, purifying selection must be maintaining protein interactome resilience in the LTEE. In addition, I find a general trend that loss-of-function mutations *increase* network resilience in REL606. This finding shows that positive selection has not optimized the interactome resilience of REL606: indeed, the interactome resilience of REL606 appears to be closer to a local minimum than to a local maximum.

Protein interactome resilience could be maintained by direct selection, or as a byproduct of selection on phenotypes that correlate with interactome resilience. In previous work, I found evidence for purifying selection on essential genes in metagenomic time series covering 60,000 generations of the LTEE (Maddamsetti and Grant 2020b), as well as evidence of purifying selection on highly interacting genes (Maddamsetti 2020). Consistent with those findings, the LTEE genomes analyzed here show evidence of purifying selection on essential genes, which are also highly enriched for protein-protein interactions. I then asked whether purifying selection on essential genes could explain the maintenance of interactome resilience in the LTEE. My analyses suggest that this is not the case: disruptions of essential genes are qualitatively similar to disruptions of non-essential genes, in regard to their effects on network resilience (Figure 4 and Supplementary Figure S1). These findings still leave open the broader question of whether protein interactome resilience is under direct selection, or is a byproduct of selection on other, unknown, correlated phenotypes. In this vein, it would be interesting to ask whether variation in interactome resilience correlates with evolvability in the sense of the distribution of fitness effects (DFE) for beneficial mutations (Mustonen and Lässig 2010; Woods, et al. 2011; Łuksza and Lässig 2014; Levy, et al. 2015; Ba, et al. 2019), or with mutational robustness in the sense of the DFE for deleterious mutations (Johnson, et al. 2019).

This work has important limitations. First, the resampling procedure used to generate randomized networks does not maintain the block structure of large deletion mutations. For this reason, I analyzed the protein interaction degree and genomic interaction distance distribution of genes affected by large deletions in the LTEE. This analysis did not uncover any systematic biases that would affect the broad import of my findings. Second, the resampling procedure does not preserve the phylogenetic structure within each population (i.e. randomized networks at later time points are not subnetworks of the randomized networks at earlier timepoints), for the sake of computational tractability. Duplication and amplification mutations are also ignored, owing to their rarity in these data (Tenaillon, et al. 2016), and the evolution of new interactions is ignored due to a lack of data. Third, it is possible that gene essentiality evolves in the LTEE. Even though purifying selection on essential genes in the ancestral clone does not appear to be sufficient to cause selection for network resilience, it is possible that essential genes in the evolved clones make a greater contribution to network resilience than essential genes in the ancestral clone. Finally, the LTEE was specifically designed to minimize ecological complexity (Lenski et al, 1991). Given the significant correlation between network resilience and ecological complexity reported by Zitnik et al. (2019), it is possible that network resilience may often evolve under positive selection in nature, but not in the controlled and largely constant abiotic conditions of the LTEE.

Finally, there is an intriguing connection between network resilience and the deterministic mutation hypothesis for the evolution of sex (Kondrashov 1988; Azevedo, et al. 2006). Loss-of-function mutations may have little effect on network integrity when they occur in a genome with high network resilience. By contrast, they may have catastrophic effects on network integrity when they occur in a genome with low network resilience. The deterministic mutation hypothesis states that synergistic epistasis between deleterious mutations—such that those that together cause network fragmentation— confers a selective advantage to sex. Near a critical threshold of network resilience, additional loss-of-function mutations are more likely to fragment biological networks, which could contribute to the synergistic epistasis required by the deterministic mutation hypothesis for the evolution of sex. Gene disruptions continue to accumulate over time in each LTEE population, suggesting that it might be worthwhile to test for such synergistic interactions at a later point (Elena and Lenski 1997), especially in the context of PPI network resilience.

## Data Availability Statement

All data used is publicly available, as described in the Material and Methods. The data and analysis codes underlying this article are available on the Dryad Digital Repository (DOI pending publication). Analysis codes are also available at: https://github.com/rohanmaddamsetti/LTEE-network-analysis.

## Acknowledgements

I thank Richard Lenski, Jeffrey Barrick, and Lingchong You for guidance and advice, Tom Milledge and Duke Research Computing for research support, Nkrumah Grant for valuable discussions, and Zachary Blount for detailed comments and feedback on an earlier version of this manuscript. The LTEE is supported by a grant from the National Science Foundation (currently DEB-1951307) to Richard Lenski and Jeffrey Barrick.

---

**Supplementary Figure S1.**
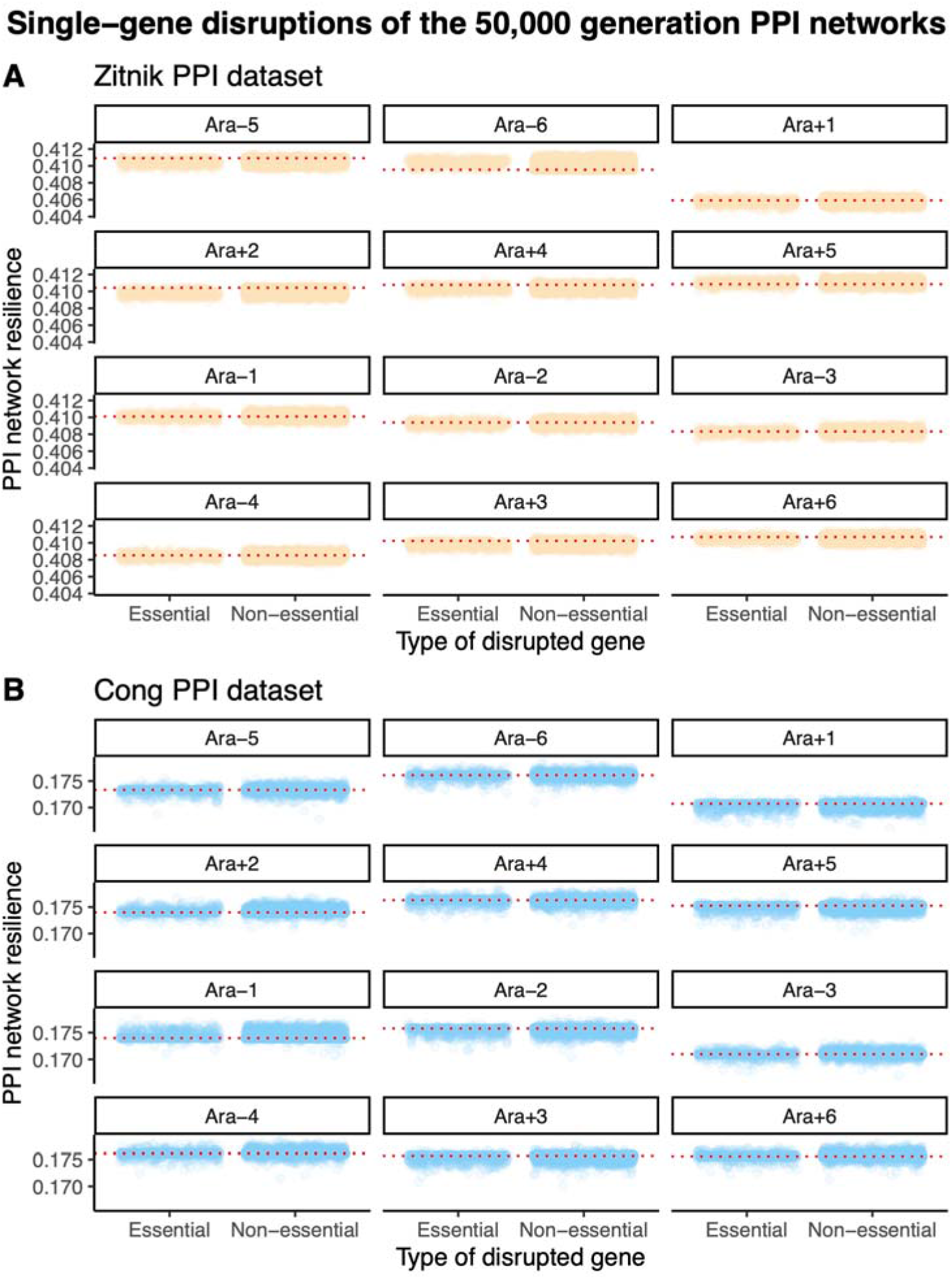
The effects of single-gene disruptions on the resilience of 50,000 generation LTEE PPI networks. Each point represents the resilience of the PPI network after a single gene has been removed. In each subfigure, the top six populations have the ancestral point-mutation rate, while the bottom six populations evolved elevated point-mutation rates. Within each population-specific panel, the distribution of resilience effects after removing essential and nearly essential genes is shown on the left, and the distribution of resilience effects after removing non-essential genes is shown on the right. The dashed red line in each population-specific panel indicates the resilience of the original network for the given 50,000 generation LTEE clone. A) Analysis based on the *E. coli* PPI network published in Zitnik et al. (2019). B) Analysis based on the *E. coli* PPI network published in Cong et al. (2019).

